# Landmark-based geometric morphometrics: Influential landmarks

**DOI:** 10.1101/2025.07.14.664645

**Authors:** Dujardin Jean-Pierre, Sriwichai Patchara, Samung Yudthana, Ruangsittichai Jiraporn, Sumruayphol Suchada, Dujardin Sebastien

## Abstract

Geometric morphometrics based on two-dimensional landmarks is a powerful tool for distinguishing morphologically similar or cryptic taxa, an important asset in the fight against medically and veterinary important arthropods. While it is commonly assumed that increasing the number of landmarks should improve discriminatory power by capturing more shape information, our findings challenge this assumption. In terms of shape discrimination, we demonstrate that small subsets of landmarks can outperform full sets of landmarks. Examples are given in 6 insect families: Culicidae, Glossinidae, Muscidae, Psychodidae, Reduviidae and Tabanidae. In all of these examples where landmark-based geometric morphometry was effective in separating morphologically close taxa, the total number of landmarks was not as effective as some significantly smaller subsets. To find such performing subsets, we used a random approach. Thus, for each number of landmarks (subsets), we examined a random sample of their many possible combinations. This random search was compared to a simpler approach, called the hierarchical method, based on the contribution of each landmark to the overall distance between shapes. Both procedures have been integrated into the XYOM online software, providing accessible tools for efficient landmark selection and improved morphometric analysis.

**Author summary:** Landmark-based geometric morphometrics describes shape in direct relation to the number of landmarks used. It is commonly assumed that increasing the number of landmarks allows for more information about shape, and when discriminating between groups or taxa, this strategy is expected to improve classification accuracy. Our results challenge this assumption. We demonstrate that subsets of landmarks, as small as three or four, can outperform the species classification obtained by the full set of landmarks, and we propose two methods for identifying them. We analyze the possible causes of these counter-intuitive results and the perspectives they could open for morphometric studies.

## Introduction

Species determination based on morphology is generally satisfactory but can become tricky for closely related or cryptic species, requiring a high level of entomological knowledge and experience. Due to some limitations of the morphological approach in taxonomy, such as the fragility of diagnostic characters, or the absence of an expert in a rare group of arthropods, complementary or alternative techniques have been developed, genetic, molecular, physical, or morphometric ones. Our study concerns morphometry, and in particular its most recent development: landmark-based, geometric morphometry [1].

This revolutionary approach captures both the size of an organ and its shape, and allows us to visualize changes in shape between individuals, even those not visible to the naked eye. Today, modern morphometrics may be seen as a rapid and cost-effective complementary tool, or sometimes an alternative means, to identify cryptic species of arthropods that often require molecular machinery to be distinguished [2].

To improve the ability of shape variables to correctly distinguish organisms, it seems logical to capture as much detail as possible. In the case of an insect wing or any other relevant body part, this often means digitizing more landmarks.

A larger number of landmarks may result in too many shape variables to describe few individuals. As an acceptable solution to possible multidimensionality problems, a subset of few first PCs of the variables has commonly been used. It reduces the number of final shape variables and extracts the main information [3, 4].

As far as we know, only one study considered the idea to use less landmarks, only the important ones. Using Monte Carlo simulations of forms designed with 5 landmarks, [5] discussed criteria based on the eigenvalues of their correlation matrix, suggesting that some landmarks are probably redundant, or bring little additional information.

Using published data, we tried to answer the following question: to reclassify taxa using wing landmarks geometry, is it more effective to select only some most informative landmarks rather than use all available ones?

Our findings demonstrate that smaller subsets of carefully chosen landmarks can provide better taxonomic discrimination than full landmark sets. We provisionally explain this counter-intuitive result both by the particular nature of the Procrustes superposition, by the uneven quality of the landmarks and by the variety of parameters which act on the conformation of organisms.

## Materials and Methods

### Culicidae

#### Aedes

The mosquitoes belonging to the genus *Aedes* Meigen, 1818 are vectors of viruses responsible for severe diseases, among which yellow fever, dengue, Chikungunya and Zika. *Aedes aegypti* (Linnaeus, 1762), *Ae. albopictus* (Skuse, 1894) and *Ae. scutellaris* (Walker, 1858) examined here were collected in Thailand. All of these groups were satisfactorily distinguished at 16 wing landmarks by [6], except for the female pair *Ae. scutellaris* and *Ae. albopictus*. Using the full set of the 16 landmarks used by [6] and a number of subsets going from 3 to 15 landmarks, we performed here again a separate study of males and females, comparing *Ae. albopictus* with *Ae. scutellaris, Ae. aegypti* with *Ae. scutellaris*, and *Ae. albopictus* with *Ae. aegypti*.

#### Anopheles

The *Anopheles* Meigen, 1818 mosquitoes are the vectors of paludism, one of the world’s deadliest parasitic diseases. The coordinates of female *Anopheles minimus* Theobald, 1901 and *An. harrisoni* Harbach & Manguin, 2007 were those used by [7]. We used the data comparing 67 female *An. minimus* and 22 *An. harrisoni*.They are based on the same number of landmarks (16) as for the *Aedes* species (see), and collected in the same order. We then explored the reclassification power of less landmarks.

Two group sizes were also compared between *Anopheles sawadwongporni* Rattanarithikul & Green, 1987 and *An. pseudowillmori* (Theobald, 1910). In this comparison the raw coordinates corresponded to 17 landmarks as in [8]. Here, different operators, T and J (one of us), used a different sampling of the mosquitoes: 41,36 and 26,28, respectively. These two sampling were extracted from the same data [8].

### Glossinidae

The tsetse flies are responsible for the transmission of trypanosomiasis to the human (sleeping disease) and to the livestock (nagana). We used two subspecies of *Glossina palpalis* from Ivory Coast (Côte d’Ivoire): *Glossina palpalis gambiensis* (*Gpg*) and *Glossina palpalis palpalis* (*Gpp*) [9].

In the original publication ( [9]), the 11 landmarks were annotated by two operators: B(erte) and TA(Ta) and no comparison was perfomed between them. The samples digitized by B were composed of 40 *Gpg* and 18 *Gpp*. The sample digitized by TA were the 40 same *Gpg* and 24 different *Gpp* coming from another geographic location of southern Ivory Coast (Côte d’Ivoire). Thus, using the 40 *Gpg* sample we could compare the two operators (B and TA) using either the complete set of landmarks or a number of subsets of them.

### Muscidae

The species belonging to the *Stomoxys* Geoffroy, 1972 genus are hematophagous flies of veterinary and medical importance, acting as vectors of many pathogens. They are also a major irritant pest of both livestock and wildlife, and a nuisance for humans [10]. To explore the discriminatory power of few landmarks, we used the wing coordinates of a maximum of 10 landmarks as digitized by [11] to compare male *Stomoxys pullus* Austen, 1909 and *Stomoxys uruma* Shinonaga et Kano, 1966.

### Reduviidae

The North American *Triatoma protracta* (Uhler) (1894) is a potential vector of Chagas disease in North America. It has been described as a complex of various subspecies [12]. We included two of them in our study: *T. p. woodi* and *T. p. protracta*. We used the data corresponding to 13 landmarks as published by [13].

### Tabanidae

Tabanidae are hematophagous flies of veterinary and medical importance, biological and mechanical vectors of many parasites [14]. From the publication of [15] we used the coordinates at 22 landmarks and at the many subsamples of them to compare *Tabanus megalops* Walker, 1854 and *Tabanus striatus* Fabricius, 1787.

### Psychodidae

*Sergentomyia hodgsoni* (Sinton, 1933) and *Phlebotomus stantoni* (Newstead 1914), as different genera, are not to be considered as morphologically close species, and they were previously shown to be perfectly separated by 12 landmarks of the wings [16]. They are used in the present study to examine the fate of a perfect taxonomic signal when the number of landmarks is progressively reduced.

### Qualifying landmarks

Using two species from each family, we compared the reclassification accuracy obtained with the total number of landmarks and the ones obtained with successively lower numbers of landmarks. Thus, we built different subsets of landmarks, decreasing from the total set of landmarks up to a minimum subset of 3 landmarks.

For each subset of landmarks, we performed the Procrustes superposition (GPA) and used an estimate of the Procrustes distance to re-classify the compared groups.

To give each landmark (or each subset of landmarks) an estimation of its contribution to the discrimination of two groups, two main methods were applied. The first one, called the “random search method”, was based on the maximum performance in a given subset of landmarks among a sample of random combinations of landmarks (see next section). The second one, that we call the “hierarchical method”, was based on the contribution of each landmark to the global shape distance between two configurations after aligning them using the complete set of landmarks.

### The random search method (“Ra”)

The first method explored at least two times a random sample (N=90) from the many possible combinations of landmarks configurations, going from 3 landmarks to the total used by each cited reference ( [6], [7], [8], [11], [16], [15], [9], [13]).

The number of different combinations of landmarks that could be assembled to form a given subset of k landmarks was computed as a classical combination (not arrangements), as:

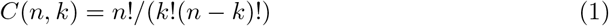

where: *n* is the complete number of landmarks, *k* is the subset of landmarks.

We computed the correlation between the allometric residue of shape for each Ra subset and the accuracy score it allowed, as well as the correlation between the accuracy scores of the subsets and the Procrustes distance.

The reclassification scores obtained by the successive subset of *k* landmarks allowed us to distinguish successful and unsuccessful configurations. We considered a subset as a successful one when its reclassification accuracy was equivalent or higher than the one obtained with the complete set of landmarks.

### The hierarchical method

This simple method performed a previous hierarchy of landmarks after alignment of forms (and projection onto a tangent plane), based on the Euclidean distances between pairs of homologous landmarks. To select influential landmarks for each subsets, we followed the order of them sorted according to their respective contribution to the global distance between shapes. To account for potential bias introduced by the least-square (LS) optimization alignment, we also performed the alignment using the resistant-fit (RF) method [17]. In addition to be computationally simple, the hierarchy of landmarks could be guessed visually from the graphical superposition of configurations (Fig 1).

**Fig 1.**
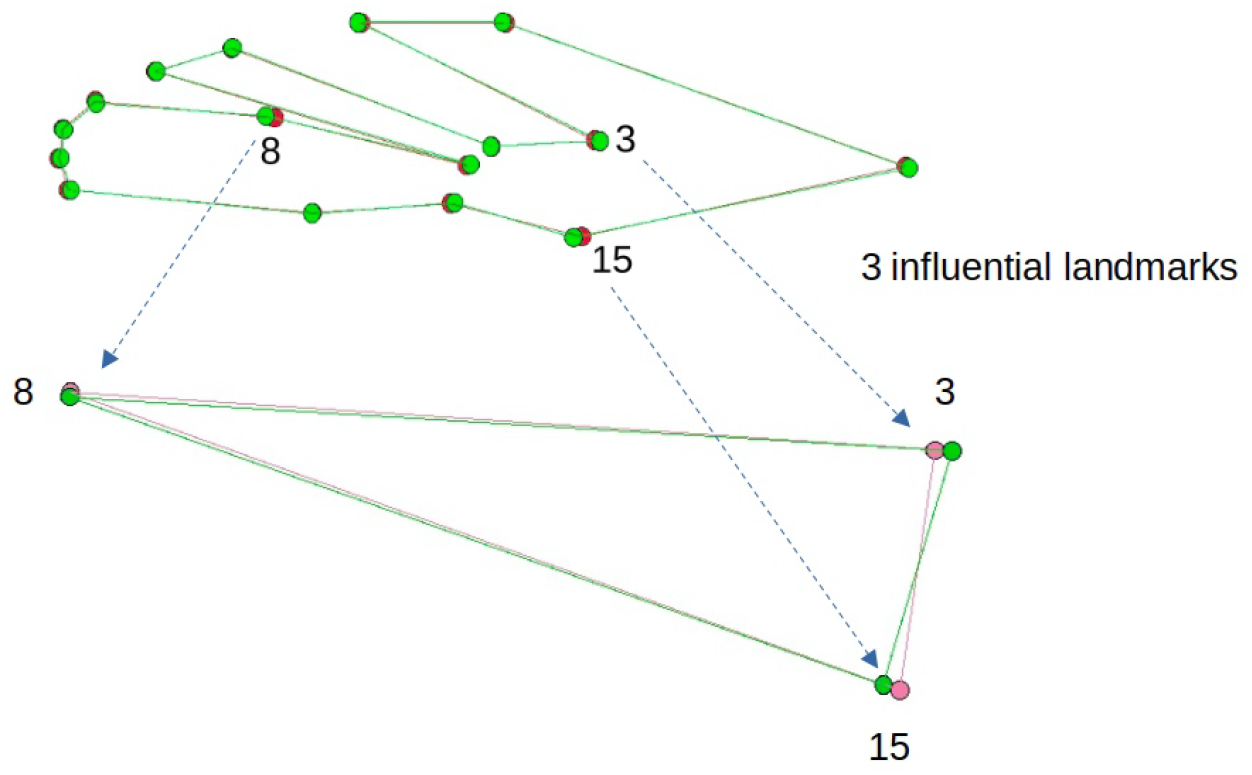
**Procrustes superposition of 16 landmarks of *Anopheles harrisoni* and *An. minimus* [7]. Three influential landmarks may be visually identified**

### Classification tools

The reclassification of the samples was based on shape variables, not on size. Shape variables were computed according to the generalized Procrustes analysis (GPA) [18], and their residual allometry was computed for subsequent correlation analyses.

To explore the ability of each subset of landmarks to recognize the correct species, we used the Kmeans approach with K the number of compared species or groups [19]. The centroid initialization method was based on naive sharding (https://www.kdnuggets.com/2017/03/naive-sharding-centroid-initialization-method.html). We used the Euclidean distance between principal components of shape, assumed to be highly correlated to the non-Euclidean Procrustes distance itself when differences between shapes are small [20, 21]. The percentage of individual matches between the groups computed by the Kmeans method and the pre-established ones represents here the reclassification score.

### Software

All computation were performed using specific scripts derived from the XYOM (version III) functions. They are proposed in the miscellaneous section of the online XYOM software (https://xyom.io). The resistant-fit alignment procedure was performed using the R “opSup” function from [21].

## Results

We performed reclassification studies on six families of insects: Culicidae, Glossinidae, Muscidae, Psychodidae, Reduvidae and Tabanidae. In our Tables, the naming of the landmarks are the order numbers as found in the published papers (see Table 1).

**Table 1.**
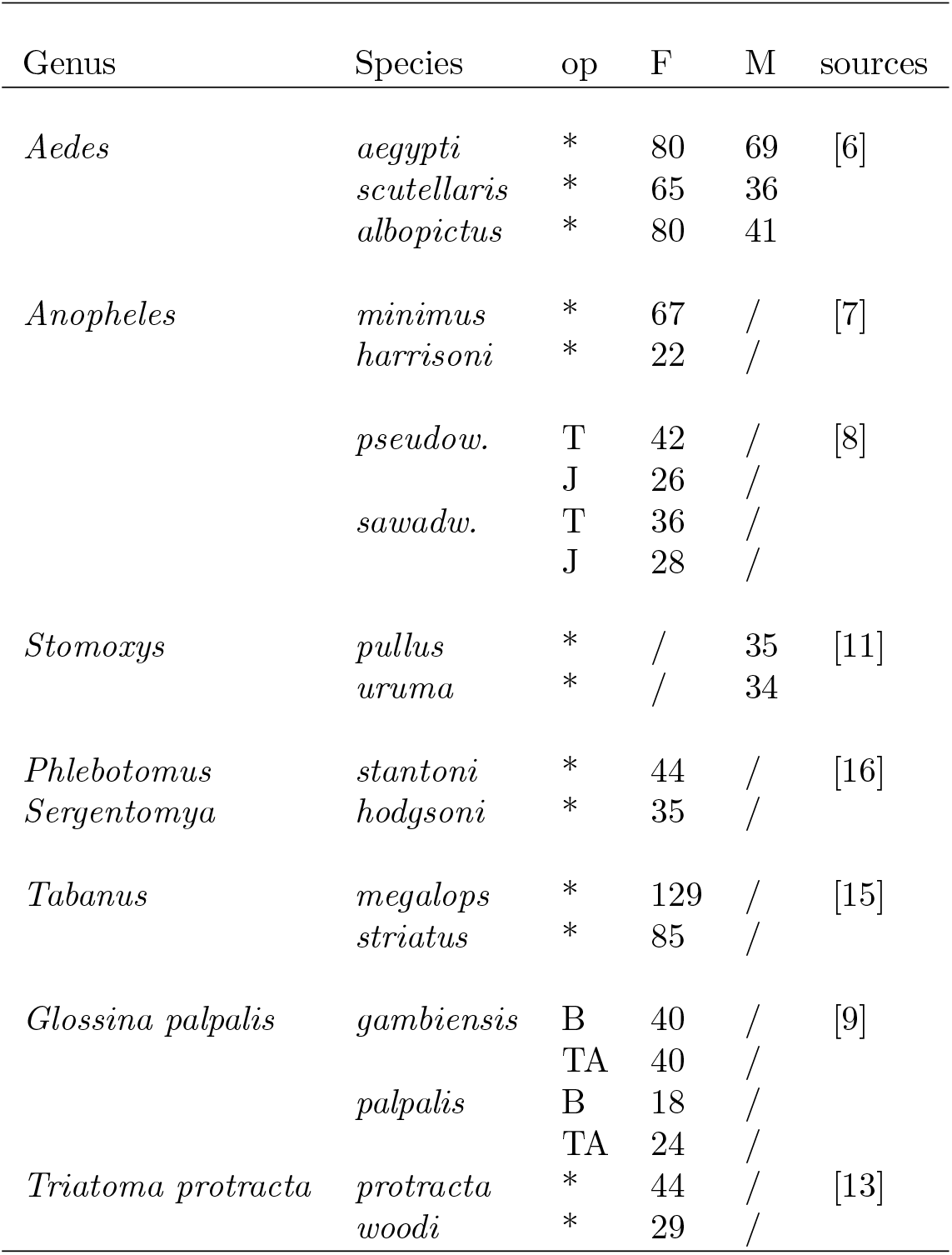
Materials. Genera and Species. *, not mentioned; op, operator; F, females; M, males; *peudow*., *pseudowillmori*; *sawad*., *sawadwongporni*.

| Genus | Species | op | F | M | sources |
| --- | --- | --- | --- | --- | --- |
| *Aedes* | *aegypti* | * | 80 | 69 | [6] |
|  | *scutellaris* | * | 65 | 36 |  |
|  | *albopictus* | * | 80 | 41 |  |
| *Anopheles* | *minimus* | * | 67 | / | [7] |
|  | *harrisoni* | * | 22 | / |  |
|  | *pseudow.* | T | 42 | / | [8] |
|  |  | J | 26 | / |  |
|  | *sawadw.* | T | 36 | / |  |
|  |  | J | 28 | / |  |
| *Stomoxys* | *pullus* | * | / | 35 | [11] |
|  | *uruma* | * | / | 34 |  |
| *Phlebotomus* | *stantoni* | * | 44 | / | [16] |
| *Sergentomya* | *hodgsoni* | * | 35 | / |  |
| *Tabanus* | *megalops* | * | 129 | / | [15] |
|  | *striatus* | * | 85 | / |  |
| *Glossina palpalis* | *gambiensis* | B | 40 | / | [9] |
|  |  | TA | 40 | / |  |
|  | *palpalis* | B | 18 | / |  |
|  |  | TA | 24 | / |  |
| *Triatoma protracta* | *protracta* | * | 44 | / | [13] |
|  | *woodi* | * | 29 | / |  |

Table 2 shows the total number of subsets randomly generated (column “n”) for each pair of taxa, ranging from 505 (in *Stomoxys*) to 1642 (in *Tabanus*). From this total, only a fraction provided similar or better reclassification accuracy scores than the one obtained with the complete set of landmarks: this proportion is shown in the same Table (column “successful”). This table 2 also shows some statistics related to the Ra method. For the many reclassification analyses (from 505 to 1642) performed at each species comparison, the linear correlation was computed between the accuracy and either the Procrustes distance or the allometric residue of shape variables. As expected, the correlation between accuracy and distance between shapes was always positive, although not always highly positive. For *Glossina, Stomoxys* and Psychodidae, the accuracy scores clearly increased with the allometric residue, while for *Triatoma, Tabanus* and some Culicidae, higher accuracy was obtained with less allometric effect. For each comparison between two taxa, the random search method allowed to find the most performant subset of landmarks for an optimal reclassification (see Table 3).

**Table 2.** Correlation coefficient between the reclassification accuracy scores of landmarks subsets and (i) the allometric residues of shape variables (column “Acc-All”), as well as (ii) the Procrustes distances between the taxa (column “Acc-Pro”. n, number of subsets examined; successful, the proportion of the n subsets that provided an accuracy equal or higher than the one of the complete set of landmarks; sa, *sawadwongporni*; ps, *pseudowillmori*; mi, *minimus*; ha, *harrisoni*; ae, *aegypti*; al, *albopictus*; sc, *scutellaris*; pa, *palpalis*; ga, *gambiensis*; pr, *protracta*, wo, *woodi*; me, *merus*; st, *striatus*; pu, *pullus*; ur, *uruma*; Ph, *Phlebotomus*; Se, *Sergentomyia*.

|  | Acc-All | Acc-Pro | n | successful |
| --- | --- | --- | --- | --- |
| *Anopheles* |  |  |  |  |
| sa/ps 26/28 | 0.18 | 0.52 | 1187 | 40% |
| sa/ps 42/36 | 0.04 | 0.55 | 1187 | 55% |
| mi/ha 67/22 | -0.52 | 0.39 | 1096 | 51% |
| *Aedes* |  |  |  |  |
| ae/al F 80/80 | -0.40 | 0.36 | 1096 | 12% |
| ae/al M 69/41 | 0.43 | 0.26 | 1096 | 14% |
| ae/sc F 80/65 | 0.15 | 0.36 | 1096 | 25% |
| ae/sc M 69/36 | -0.22 | 0.29 | 1096 | 17% |
| al/sc F 80/65 | 0.13 | 0.36 | 1096 | 67% |
| al/sc M 41/36 | 0.21 | 0.32 | 1096 | 18% |
| *Glossina* |  |  |  |  |
| ga/pa1 40/18 | 0.91 | 0.54 | 606 | 37% |
| ga/pa2 40/24 | 0.86 | 0.64 | 606 | 10% |
| *Triatoma* |  |  |  |  |
| pr/wo 44/29 | -0.25 | 0.35 | 811 | 34% |
| *Stomoxys* |  |  |  |  |
| pu/ur 35/34 | 0.91 | 0.55 | 505 | 31% |
| *Tabanus* |  |  |  |  |
| me/st 129/85 | -0.62 | 0.75 | 1642 | 40% |
| Sandflies |  |  |  |  |
| Ph/Se 44/35 | 0.83 | 0.62 | 708 | 54% |

**Table 3.**
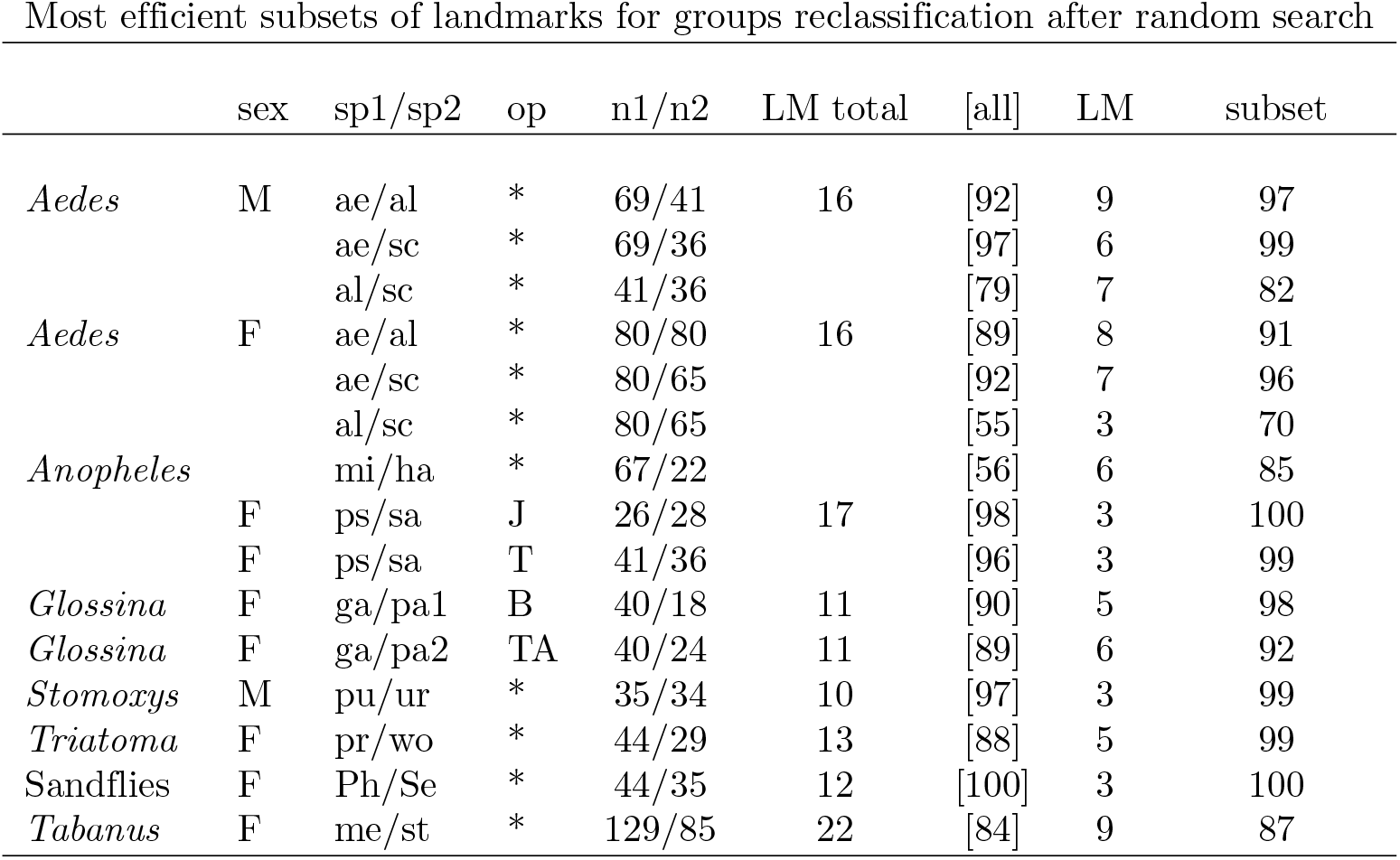
Most efficient subsets of landmarks for groups reclassification based on Kmeans after random search (Ra). [all], accuracy computed from the total number of landmarks; subset, accuracy computed from the subset of landmarks which produced the best reclassification score: the number of landmaks of these subsets is given in column “LM”. sa, *sawadwongporni*; ps, *pseudowillmori*; mi, *minimus*; ha, *harrisoni*; ae, *aegypti*; a, *albopictus*; s, *scutellaris*; pa, *palpalis*; ga, *gambiensis*; pr, *protracta*, wo, *woodi*; me, *merus*; st, *striatus*; pu, *pullus*; ur, *uruma*; Ph, *Phlebotomus*; Se, *Sergentomyia*

### Interspecific comparisons

In addition to the Table 2 giving some statistics about the Ra method, the main information provided by our study may be summarized by considering the subsets of only three and four influential landmarks (Tables 4, 5).

**Table 4.** Most efficient reclassification triangles of landmarks after least-square alignment (LS), resistant-fit alignment (RF) or after random search (Ra). The corresponding computed reclassification accuracies are in the columns [LS], [RF] and [Ra]. Rightmost column [all], accuracy computed from the total number of landmarks. sa, *sawadwongporni*; ps, *pseudowillmori*; mi, *minimus*; ha, *harrisoni*; ae, *aegypti*; a, *albopictus*; s, *scutellaris*; pa1, *palpalis*; pa2, *palpalis*; ga, *gambiensis*; pr, *protracta*, wo, *woodi*; me, *merus*; st, *striatus*; pu, *pullus*; ur, *uruma*; Ph,*Phlebotomus*; Se,*Sergentomyia*

| Most discriminating triangles of landmarks |  |  |  |  |  |  |  |  |
| --- | --- | --- | --- | --- | --- | --- | --- | --- |
| Species n1/n2 | op | LS | RF | Ra | [LS] | [RF] | [Ra] | [all] |
| *Anopheles* |  |  |  |  |  |  |  |  |
| sa/ps 26/28 | J | 4,13,12 | 4,11,13 | 2,4,17 | 59 | 94 | 98 | [98] |
| sa/ps 42/36 | T | 4,13,12 | 13,12,11 | 2,4,17 | 67 | 60 | 100 | [98] |
| mi/ha 67/22 | * | 15,8,3 | 15,8,3 | 3,8,15 | 82 | 82 | 82 | [56] |
| *Aedes* |  |  |  |  |  |  |  |  |
| ae/a F 80/80 | * | 7,4,16 | 7,16,4 | 1,2,7 | 61 | 61 | 89 | [89] |
| ae/a M 69/41 | * | 7,4,2 | 7,4,16 | 7,14,15 | 60 | 60 | 89 | [92] |
| ae/s F 80/65 | * | 7,4,5 | 7,4,5 | 7,14,5 | 82 | 82 | 93 | [92] |
| ae/s M 69/36 | * | 7,4,5 | 16,7,4 | 7,14,16 | 72 | 63 | 98 | [97] |
| al/s F 80/65 | * | 13,15,4 | 13,16,15 | 6,8,13 | 60 | 54 | 70 | [55] |
| al/s M 41/36 | * | 5,1,14 | 5,16,1 | 1,2,16 | 78 | 79 | 79 | [79] |
| *Glossina* |  |  |  |  |  |  |  |  |
| ga/pa1 F 40/18 | B | 1,9,5 | 9,5,10 | 2,6,9 | 84 | 59 | 88 | [90] |
| ga/pa2 F 40/24 | TA | 2,10,6 | 6,4,10 | 2,6,9 | 73 | 48 | 75 | [89] |
| *Triatoma* |  |  |  |  |  |  |  |  |
| pr/wo 44/29 | * | 5,4,3 | 5,3,4 | 3,6,12 | 45 | 45 | 96 | [88] |
| *Stomoxys* |  |  |  |  |  |  |  |  |
| pu/ur 35/34 | * | 1,7,2 | 1,7,2 | 1,2,4 | 87 | 87 | 99 | [97] |
| *Tabanus* |  |  |  |  |  |  |  |  |
| me/st 129/85 | * | 2,4,22 | 2,4,3 | 2,17,21 | 83 | 74 | 86 | [84] |
| Sandflies |  |  |  |  |  |  |  |  |
| Ph/Se 44/35 | * | 1,12,2 | 1,12,2 | 3,11,12 | 95 | 95 | 100 | [100] |

**Table 5.** Most efficient reclassification squares of landmarks after least-square alignment (LS), resistant-fit alignment (RF) or after random search (Ra). The corresponding computed reclassification accuracies are in the columns [LS], [RF] and [Ra]. Rightmost column [all], accuracy computed from the total number of landmarks. sa, *sawadwongporni*; ps, *pseudowillmori*; mi, *minimus*; ha, *harrisoni*; ae, *aegypti*; al, *albopictus*; sc, *scutellaris*; pa1, *palpalis*; pa2, *palpalis*; ga, *gambiensis*; pr, *protracta*, wo, *woodi*; me, *merus*; st, *striatus*; pu, *pullus*; ur, *uruma*; Ph, *Phlebotomus*; Se, *Sergentomyia*. For sample sizes, please refer to the previous Table 4.

| Most discriminating squares of landmarks |  |  |  |  |  |  |  |  |
| --- | --- | --- | --- | --- | --- | --- | --- | --- |
| Species n1/n2 | op | LS | RF | Ra | [LS] | [RF] | [Ra] | [all] |
| *Anopheles* |  |  |  |  |  |  |  |  |
| sa/ps | J | 4,13,12,11 | 4,11,13,12 | 2,3,4,17 | 95 | 95 | 100 | [98] |
| sa/ps | T | 4,13,12,17 | 4,13,12,11 | 2,3,4,17 | 95 | 96 | 100 | [98] |
| mi/ha | * | 15,8,3,7 | 15,8,3,1 | 7,8,14,15 | 76 | 58 | 83 | [56] |
| *Aedes* |  |  |  |  |  |  |  |  |
| ae/al F | * | 7,4,16,2 | 7,16,4,2 | 1,2,7,8 | 86 | 86 | 90 | [89] |
| ae/al M | * | 7,4,2,16 | 7,4,16,2 | 1,7,14,15 | 86 | 86 | 92 | [92] |
| ae/sc F | * | 7,4,5,6 | 7,4,5,6 | 1,5,7,14 | 76 | 76 | 93 | [92] |
| ae/sc M | * | 7,4,5,6 | 16,7,4,5 | 4,7,14,16 | 79 | 94 | 97 | [97] |
| al/sc F | * | 13,15,4,7 | 13,16,15,5 | 2,6,9,10 | 60 | 54 | 71 | [55] |
| al/sc M | * | 5,1,14,12 | 5,16,1,14 | 5,11,12,13 | 81 | 81 | 82 | [79] |
| *Glossina* |  |  |  |  |  |  |  |  |
| ga/pa1 F | B | 1,9,5,10 | 9,5,10,1 | 2,6,9,8 | 86 | 86 | 88 | [90] |
| ga/pa2 F | TA | 2,10,6,1 | 6,4,10,2 | 2,6,9,8 | 88 | 72 | 89 | [89] |
| *Triatoma* |  |  |  |  |  |  |  |  |
| pr/wo | * | 5,4,3,13 | 5,3,4,13 | 3,6,7,12 | 95 | 95 | 96 | [88] |
| *Stomoxys* |  |  |  |  |  |  |  |  |
| pu/ur | * | 1,7,2,5 | 1,7,2,8 | 1,2,4,10 | 97 | 99 | 99 | [97] |
| *Tabanus* |  |  |  |  |  |  |  |  |
| me/st | * | 2,4,22,3 | 2,4,3,10 | 2,17,21,22 | 83 | 80 | 86 | [84] |
| Sandflies |  |  |  |  |  |  |  |  |
| Ph/Se | * | 1,12,2,3 | 1,12,2,3 | 1,3,11,12 | 99 | 99 | 100 | [100] |

Tables 4 and 5 show the influential 3- and 4-landmark subsets, their accuracy for reclassifying the sample of two taxa, and this accuracy when using all landmarks.

Using these tables, the following main observations can be derived:

(1) The selection of LS and RF influential landmarks did not differ much. Their respective performances in reclassifying taxa suggested that, to this purpose, the least squares method was, on average, slightly better than the resistant-fit one.

(2) The most efficient Ra subsets always contained at least one landmark of the LS or RF subsets.

(3) When considering the three landmarks subsets (Table 4), the accuracy scores of the LS and RF ones could be far from the ones obtained by the Ra scores. This difference was consistently reduced for the 4 landmarks subsets (Table 5).

(4) On average, the accuracy produced by selecting LS and RF subsets of 4 landmarks (86% and 84%, respectively) was much more satisfactory than for three landmarks (73% and 69%, respectively), without however reaching the average levels of the Ra subsets of 3 (90%) or of 4 landmarks (92%).

Note that there were two interspecific comparisons showing a reclassification score compatible with chance alone: the female *Ae. albopictus* compared with the female *Ae. scutellaris* (55%), as already observed in the original publication [6], and another one, between female *Anopheles minimus* and*An. harrisoni* (56%). For the *Aedes* comparison, the best subsets could increase slightly the score, not better however than 70 %, which remained compatible with chance alone. For the *Anopheles* comparison, smaller subsets significantly improved the reclassification score, up to 85% (see Tables 3, 4 and 5).

### Interuser comparisons

Our material allowed three comparisons of specimen belonging to the same species but digitized by different operators: *Anopheles pseudowillmori, Anopheles sawadwongporni*, and *Glossina palpalis palpalis* (see Table 6). This type of comparison could be likened to a comparison between operators: the more the operators differ, the higher the expected discrimination scores.

**Table 6.** Most efficient reclassification triangles and squares of landmarks after least-square alignment (LS), resistant-fit alignment (RF) or after random search (Ra). Statistics were using either all set of landmarks (11 landmarks), or only eight of them after removing the problematic landmarks LM 1, LM2 and LM11. The corresponding reclassification accuracies are in the columns [LS], [RF] and [Ra]. Rightmost column [all], accuracy computed from the total number of landmarks (11, and 8). The order number of the landmarks is the one related to the total number (11) of landmarks. ga/pa1 40 *gambiensis* compared with 18 *palpalis* by one operator (Berte) and ga/pa2 40/24, the 40 same *gambiensis* individuals compared with another, geographically distant, sample of 24 *palpalis* annotated by another operator (TA); F, females;

| Removing problematic landmarks to identify true influential landmarks |  |  |  |  |  |  |  |  |
| --- | --- | --- | --- | --- | --- | --- | --- | --- |
| *Glossina* | op | LS | RF | Ra | [LS] | [RF] | [Ra] | [all] |
| 11 landmarks |  |  |  |  |  |  |  |  |
|  |  | Triangles |  |  |  |  |  |  |
| ga/pa1 | B | 1,9,5 | 9,5,10 | 2,6,9 | 84 | 59 | 88 | [90] |
| ga/pa2 | TA | 2,10,6 | 6,4,10 | 2,6,9 | 73 | 48 | 75 | [89] |
|  |  | Squares |  |  |  |  |  |  |
| ga/pa1 | B | 1,9,5,10 | 9,5,10,1 | 2,6,9,8 | 86 | 86 | 88 |  |
| ga/pa2 | TA | 2,10,6,1 | 6,4,10,2 | 2,6,9,8 | 88 | 72 | 89 |  |
| 8 landmarks |  |  |  |  |  |  |  |  |
|  |  | Triangles |  |  |  |  |  |  |
| ga/pa1 | B | 9,5,10 | 9,5,10 | 5,6,9 | 59 | 59 | 95 | [91] |
| ga/pa2 | TA | 10,5,9 | 9,8,10 | 5,6,8 | 52 | 64 | 83 | [81] |
|  |  | Squares |  |  |  |  |  |  |
| ga/pa1 | B | 9,5,10,6 | 9,5,10,6 | 5,6,9,10 | 95 | 95 | 95 |  |
| ga/pa2 | TA | 10,5,9,6 | 9,8,10,5 | 5,6,8,4 | 84 | 59 | 83 |  |

Based on all landmarks, the comparison of the *Anopheles* conspecific samples annotated by different users (T and J) produced reclassification scores in line with chance expectations (52%, 54%), weakly improved by more discriminant subsets of influential landmarks (69% and 71%).

The conspecific comparison of 18 and 24 *Glossina p. palpalis* could not be reduced to a simple inter-user comparison because these samples came from different geographical areas: they therefore included possible differences due to local geographical effects on shape. On the contrary, the comparison of *G. p. gambiensis* between two operators was indeed a simple inter-operator comparison, because the specimens (and their images) were exactly the same. Despite this, the reclassification score (with all landmarks) was surprisingly high (91%). Since both operators scanned exactly the same images, influential landmarks indicated inter-operator differences, and only that. The problematic landmarks were identified as LM1, LM2, and LM11 (by both methods, details not shown). Their removal lowered the expected reclassification score to values compatible with two groups composed of the same specimens (56%), a score that no subset of landmarks could significantly improve. Furthermore, these significant differences between users probably explain why each operator generated different influential landmarks when comparing the subspecies *G. p. gambiensis* and *G. p. palpalis*. Indeed, after removing the problematic landmarks LM1, LM2 and LM11, the landmarks contributing to the divergence of the subspecies became the same for both operators (see table 6).

## Discussion

Our results were not intended to compare different insect species or groups, but rather to select the most efficient landmark configuration to discriminate between them. The number of landmarks required for optimal discrimination between taxa should not be confused with the minimum number of landmarks required to characterize shape and size variation [22]. Thus, to select the most efficient subset of landmarks, we used two different strategies, both requiring at least one prior Procrustes superposition. The efficiency of both methods was estimated by the subsequent reclassification accuracy scores.

The first strategy (Ra) was to perform a random search for smaller coordinate system configurations that would be as effective, if not more effective, than the initial configurations in reclassifying taxa. This approach required performing a very large number of Procrustes superpositions and subsequent reclassifications (Table 2). It is laborious, but close to an exhaustive search, and able to suggest more than one solution.

The second method (LS, RF) relies on the hierarchy of contributions of landmarks to the overall shape-based distance between two configurations. It is less efficient than the first but it is fast and relies on intuitive logic.

### Selecting influential landmarks

(1) Random search method (Ra)

In this approach, the relevant composition of the landmark subsets was selected after performing the Kmeans reclassification process, not before. Due to obvious technical limitations, we did not use the full set of possibilities but a sample of them, in fact a maximum of 90 combinations per subset, where sometimes tens of thousands or more were possible. Due to this random sampling step, the identity of influential landmarks could vary slightly from one run to another.

(2) Hierarchical method (LS and RF).

This method was entirely based on the Procrustes alignment method, before the reclassification process. We compared the least squares (LS) alignment method and the resistant fitting (RF) one. The LS method (commonly known as GPA) has the disadvantage of attributing the shape change to all landmarks rather than to the truly influential landmarks. Two different configurations on the same landmark would produce a superposition suggesting differences on other landmarks as well (the “Pinocchio effect”). The resistant fitting (RF) optimization procedure was developed to mitigate this disadvantage. In terms of reclassification accuracy, the LS-subsets of 4 landmarks was, on average, much more satisfactory than for 3 landmarks (86% versus 73%, respectively). Unexpectedly, the LS-subsets showed slightly better reclassification scores than the RF-subsets.

In terms of influential landmarks identity, our data did not reveal any significant difference between the two strategies, the random search strategy and the hierarchical one. In six comparisons, the same three most influential landmarks were found; in the remaining ten comparisons, two of the three most influential landmarks were found in both methods (Table 4).

However, the accuracy scores obtained by the landmarks subsets as suggested by the hierarchical approach was generally lower than the ones suggested by the random search (Ra). This might be explained by the fact that the most contributing landmarks set by the hierarchical method are initially computed for all landmarks, once for all. Their influential effect might be different when combined with another set of landmarks, as in smaller configurations (detailed results not shown).

### Classification method

To reclassify two taxa, we used an unsupervised, non-hierarchical agglomerative clustering analysis, the Kmeans method, instead of the frequently used linear discriminant analysis (LDA), which is a supervised method.

The Kmeans algorithm compares individuals and mobile centroids to suggest possible natural groupings, while the LDA compares pre-established groups. As a consequence, the LDA, although powerful, is subject to some statistical limitations [23]. Moreover, from a morphometrician point of view, the Mahalanobis distance (the metric of LDA) has the inconvenient to “distort” the Procrustes geometry [24].

The Kmeans method used the Euclidean distances between principal components of shape variables, thus after LS alignment and orthogonal projection onto a tangent plane (GPA). These Euclidean distances are assumed to be highly correlated to the non-Euclidean Procrustes distance itself when differences between shapes are small [20, 21]. The use of the naive-sharding Kmeans initialization method was adopted to tentatively reduce the inconvenient of random selection of K initial centroids, which is liable to induce instability of the final output. Thus, because of the Kmeans algorithm, the re-classification accuracy was not depending only on the global distance between two shapes, but on the individual Euclidean distances to the mobile centroids of other clusters of individuals. This algorithm explains why the correlation coefficient between accuracy scores and Procrustes distances between species was not as high as might be expected (Table 2).

### Possible causes of improved performance with fewer landmarks

The beneficial effect of using some smaller configuration of landmarks is reminiscent of the well-known phenomenon called “the curse of multidimensionality” [25]. With more parameters, models can become overly complex, fitting the noise rather than the real signal.

In our study, however, the number of variables was not particularly high, remaining well below the sample sizes, and yet our results suggested that reducing the dimensionality of the data could be beneficial.

Various reasons related to the morphometric method itself could also explain the higher discrimination of some reduced subsets of landmarks.

(i) The first one that comes to mind would simply be the nature of the distance used to perform the classifications: (an estimate of) the Procrustes distance. Because it is not divided by the number of landmarks, it does not depend on their number, but on how those landmarks influence the overall shape. Adding or removing landmarks only affects the distance if this operation contributes to global alignment differences.

(ii) Another possible cause is that our results are due to size effects, i.e., an increasing importance of size difference when reducing the number of landmarks. The Procrustes superposition removes the isometric change of size, not the allometric one. Does the allometric residue of shape increase with fewer landmarks? The answer is sometimes yes, sometimes no (Table 2).

There are also some other reasons, not directly related to the Procrustes superposition, why a reduced set of landmarks could perform better than the totality when comparing species.

(i) As highlighted by [5], the positional variation of some landmarks could be determined by the position of other landmarks via common developmental or biomechanical causes, and not directly related to evolutionary differences between species.

(ii) In addition to the effect of an evolutionary divergence on some landmarks positions, different environments could influence shape. Such influence has been apparent after a two-factor ANOVA on shape in an ecomorphometric study, where the influence of different environments could be eliminated by reducing the number landmarks, producing results more in accordance with species richness [26].

(iii) Finally, more artifactual circumstances could be incriminated also. One is related to the different qualities of the collected landmarks. Bookstein suggested a hierarchical graduation I, II, III [27]. Landmarks of lower quality (II,III) could introduce noise into the alignment process: removing them could leave the remaining set with sharper discriminating power. Moreover, landmarks of type I themselves are not equally shaped: some are points at the crossing of two or three narrow veins, other are similar to a small areas more than to just a point (optical microscopy). The latter may become problematic: because some landmarks are more difficult to correctly annotate, poorly experimented operators could introduce some bias at some landmarks.

### Perspectives

Between established but morphologically very close taxa, subtle differences in wing vein architecture are not random, and our data suggest that they may be represented by a very small set of landmarks, perhaps localized to specific areas of the wings (see for instance [28] for a very different approach). However, identifying influential landmarks is only informative if the possible sources of variation between two samples are well understood.

Among these sources of variation, there may be simple artifacts, as illustrated by the situation where two operators annotated exactly the same specimens. It is not recommended to use samples annotated by more than one operator, but it may be necessary to compare two operators. Our method could help to remove problematic (influential) landmarks (see Table 6). The same strategy could be used also to reduce intra-user error, improving the data for symmetry analyses.

The detection of a few influential landmarks could open new, original experimentation ideas about the way some factors affect shape. Thus, under an experimental protocol, the identification of influential landmarks could help to identify the area of the wings which is affected by a given factor, as for instance the temperature, the density, the geographic origin, etc.

If the comparison involves distinct taxa digitized by the same operator, it is acceptable to assume that the few influential landmarks are the most affected by evolutionary divergence. One might be tempted to limit the digitization effort to these landmarks alone, which would considerably reduce the workload on very large samples. However, the adoption of influential landmarks is valid if there is only a single source of variation between samples. The reality is less simple, as each species can generally undergo slight changes in form as it adapts to different environments, for example, different geographic areas. This effect is difficult to predict, and the data in our study lack examples illustrating this problem. In cases where there are multiple possible sources of variation, caution is advised before generalizing the use of influential landmarks detected in a single comparison.

## Conclusion

The blind strategy of increasing the number of landmarks to improve shape discrimination risks mixing noise and signal. Our study demonstrates that a few landmarks, sometimes as few as three, can achieve higher classification accuracy than all landmarks. Because of their importance, we refer to them as “influential landmarks”. Knowing them should help better justify the choice and number of landmarks used in morphometric studies, it could also suggest the relevant anatomical areas of global shape divergence between species.

## Acknowledgments

We thank the authors of [7], [8], [11], [15] and [9] who kindly allowed us to use their raw coordinates data to perform our analyses.

## Notes

https://drive.google.com/drive/folders/1MZ4qGHm5YO50g6nTbfdZiXY2sholCPvz

### Competing Interest Statement

The authors have declared no competing interest.

### Summary of Updates

A small change in the Abstract and a few more details in the Materials and Methods section

## References

1. Rohlf FJ, Marcus LF. A revolution in morphometrics. TREE. 1993;8(4):129–132.

2. Dujardin JP. Chapter 16 - Modern Morphometrics of Arthropods: A Phenotypic Approach to Species Recognition and Population Structure. In: Tibayrenc M, editor. Genetics and Evolution of Infectious Diseases (Third Edition). third edition ed. Elsevier; 2024. p. 385–425.

3. Krzanowski WJ. Selection of variables to preserve multivariate data structure, using principal components. Applied Statistics, 36: 22–33. 1987;.

4. Krzanowski WJ. Cross-validation in Principal Component Analysis. Biometrics, 43: 575–584. 1987;.

5. Rusakov DA. Dimension reduction and selection of landmarks. A Monte Carlo Experiment. In: Advances in Morphometrics Proceedings of the 1993 NATO-ASI on Morphometrics (Marcus LF, Corti M, Loy A, Naylor GJP and Slice D, eds) New York: Plenum Publ NATO ASI, ser A, Life Sciences pp 531–398. 1996;.

6. Sumruayphol S, Apiwathnasorn C, Ruangsittichai J, Sriwichai P, Attrapadung S, Samung Y, et al. DNA barcoding and wing morphometrics to distinguish three Aedes vectors in Thailand. Acta Tropica. 2016;159:1–10. doi:10.1016/j.actatropica.2016.03.010.

7. Chatpiyaphat K, Sumruayphol S, Dujardin J, Y S A P, Cui L, et al. Geometric morphometrics to distinguish the cryptic species Anopheles minimus and An. harrisoni in malaria hot spot villages, western Thailand. Med Vet Entomol 35(3):293–301. 2021;doi:10.1111/mve.12493.

8. Chaiphongpachara T, Sriwichai P, Samung Y, J R, Morales Vargas R, L C, et al. Geometric morphometrics approach towards discrimination of three member species of Maculatus group in Thailand. Acta Trop 192:66–74. 2019;doi:10.1016/j.actatropica.2019.01.024.

9. Kaba D, Berté D, Ta BTD, Telleria J, Solano P, Dujardin JP. The wing venation patterns to identify single tsetse flies. Infection Genetics and Evolution. 2017;47:132–139. doi:10.1016/j.meegid.2016.10.008.

10. Baldacchino F, Muenworn V, Desquesnes M, Desoli F, Charoenviriyaphap T, Duvallet G. Transmission of pathogens by Stomoxys flies (Diptera, Muscidae): a review. Parasite (Paris, France), 20, 26. 2013;doi:10.1051/parasite/2013026.

11. Changbunjong T, Sumruayphol S, Weluwanarak T, Ruangsittichai J, Dujardin JP. Landmark and outline-based geometric morphometrics analysis of three Stomoxys flies (Diptera: Muscidae). Folia parasitologica, 63. 2016;doi:10.14411/fp.2016.037.

12. Usinger RL. The Triatominae of North and Central America and the West Indies and their public health significance. Public Health Bulletin Washington. 1944;288: 81 pp.

13. Dujardin JP, Beard CB, Ryckman R. The relevance of wing geometry in entomological surveillance of Triatominae, vectors of Chagas disease. Infection Genetics and Evolution. 2007;7:161–167. doi:10.1016/j.meegid.2006.07.005.

14. Baldacchino F, Desquesnes M, Mihok S, Foil LD, Duvallet G, Jittapalapong S. Tabanids: Neglected subjects of research, but important vectors of disease agents! Infection, Genetics and Evolution. 2014;28:596–615. doi:10.1016/j.meegid.2014.03.029.

15. Changbunjong T, Prakaikowit N, Maneephan P, Kaewwiset T, Weluwanarak T, Chaiphongpachara T, et al. Landmark Data to Distinguish and Identify Morphologically close Tabanus spp. (Diptera: Tabanidae). Insects 12(11):974 doi: 103390/insects12110974. 2021;doi:10.3390/insects12110974.

16. Sumruayphol S, Chittsamart B, Polseela R, Sriwichai P, Samung Y, Apiwathnasorn C, et al. Wing geometry of Phlebotomus stantoni and Sergentomyia hodgsoni from different geographical locations in Thailand. C R Biol 340(1):37–46 Erratum in: C R Biol 2017; 340(3):195. 2017;doi:10.1016/j.crvi.2016.10.002.

17. Siegel AF, Benson RH. A robust comparison of biological shapes. Biometrics, 38(2): 341–350. 1982;.

18. Rohlf FJ. Rotational fit (Procrustes) methods. In: Rohlf F, Bookstein F, editors. Proceedings of the Michigan Morphometrics Workshop. Special Publication Number 2. The University of Michigan Museum of Zoology. Ann Arbor, MI, pp380. University of Michigan Museums, Ann Arbor; 1990. p. 227–236.

19. Pérez-Ortega J, Almanza-Ortega NN, Vega-Villalobos A, Pazos-Rangel R, Zavala-Díaz C, Martínez-Rebollar A. The ¡em¿K¡/em¿-Means Algorithm Evolution. In: Sud K, Erdogmus P, Kadry S, editors. Introduction to Data Science and Machine Learning. Rijeka: IntechOpen; 2019.

20. Rohlf FJ. Morphometric spaces, shape components and the effects of linear transformations. In: Marcus LF, Corti M, Loy A, Naylor GJP, Slice D, editors. Advances in Morphometrics. Proceedings of the 1993 NATO-ASI on Morphometrics. New York: Plenum Publ. NATO ASI, ser. A, Life Sciences; 1996. p. 117–129.

21. Claude J. Morphometrics with R. ISBN 978-0-387-77789-4, Springer Science+Business Media, LLC. 2008;330pp.

22. Watanabe A. How many landmarks are enough to characterize shape and size variation? PLoS ONE 13(6): e0198341. 2018;doi:10.1371/journal.pone.0198341.

23. Pimentel RA. An introduction to ordination, principal components analysis and discriminant analysis. p 11–28 In: RG Foottit and JT Sorensen (eds) Ordination in the study of morphology, evolution and systematics of insects: applications and quantitative genetic rationales Elsevier, New York, pp 418. 1992;.

24. Mitteröcker P, Bookstein F. Linear discrimination, ordination, and the visualization of selection gradients in modern morphometrics. Evolutionary Biology, 38(1): 100–114. 2011;.

25. Bellman R. Adaptive Control Processes: A Guided Tour. Princeton Legacy Library. Princeton University Press; 1961. Available from: https://books.google.fr/books?id=POAmAAAAMAAJ.

26. Dujardin JP, Thongsripong P, Henry AB. The Mosquito Fauna: From Metric Disparity to Species Diversity. Morphometrics, Christina Wahl (Ed), ISBN: 978-953-51-0172-7, InTech. 2012; http://www.intechopen.com/books/morphometrics/the-mosquito-fauna-from-metric-disparity-to-species-diversity.

27. Bookstein FL. Introduction to methods for landmark data. pp 216–225 In FJ Rohlf and FL Bookstein (eds), Proceedings, Michigan Morphometrics Workshop, 1988 The University of Michigan Museum of Zoology, Special Publication No 2, Ann Arbor, MI. 1990;.

28. Nolte K, Sauer FG, Baumbach J, Kollmannsberger P, Lins C, Lühken R. Robust mosquito species identification from diverse body and wing images using deep learning. Parasit Vectors 17, 372 (2024). 2024;doi:10.1186/s13071-024-06459-3.

